# Dark diversity illuminates the dim side of conservation and restoration

**DOI:** 10.1101/057315

**Authors:** Jesper Erenskjold Moeslund, Ane Kirstine Brunbjerg, Kevin Kuhlmann Clausen, Lars Dalby, Camilla Flojgaard, Anders Juel, Jonathan Lenoir

## Abstract

**1** *Dark diversity* is a promising concept for prioritizing management efforts as it focuses on missing species, i.e., species present in the regional pool, but locally absent despite suitable environmental conditions. **2** We applied the concept to a massive national plant diversity database (236,923 records from 15,160 surveys involving 564 species) to provide the first geographically comprehensive assessment of dark diversity across a large area (43,000 km^2^), at a spatial scale (~75 m^2^) relevant for conservation and restoration planning and across multiple terrestrial habitats, thus maximising its practical application potential. The likelihood for a given plant species to belong to the dark diversity pool was computed and logistically regressed against its ecological preferences (nutrient availability, pH etc.), strategies (competitor, stress tolerance, ruderal), mycorrhizal dependence and infection percentage, seed mass and maximum dispersal distance. **3** Forty-six percent of the species were absent in >95 % of the suitable sites, whereas 7 % of the species were absent in less than 60 % of sites that were deemed suitable. **4** Species that were more likely to belong to the dark diversity tended to depend on mycorrhiza, were mostly adapted to low light and nutrient levels, had poor dispersal abilities, were ruderals and had a low stress tolerance. **Synthesis and applications** Our findings have important implications for the planning and management of natural ecosystems requiring detailed knowledge of what triggers the presence/absence of individual plant species in a seemingly suitable habitat. We conclude that practitioners may need to carefully consider mycorrhizal inoculations with a suitable assemblage of fungi for certain plant species to become established. Also assisted migration might be necessary to help poor dispersers although spatial and temporal processes are also important to have in mind. Finally, it is important to vary nutrient loads making room for plant species to colonise both nutrient-poor and nutrient-rich localities.

## Introduction

Recently, Pärtel and co-workers (2011) presented a new concept coined *dark diversity*, which could prove to be a central idea for the development of effective tools for practical biodiversity management and conservation prioritization at relevant spatial scales. Dark diversity encompasses the diversity articulated by all species missing locally, even though biogeographic history and current ecological and environmental conditions suggest their presence (Pärtel, Szava-Kovats & Zobel, 2011). In other words, dark diversity is the set of species belonging to the regional species pool of a particular habitat but that are missing locally within a given site of that habitat (Pärtel, Szava-Kovats & Zobel, 2011). Note that the dark diversity concept differs from the so-called *hidden diversity* which counts species that are overlooked due to observation bias (Milberg *et al.*, 2008).

Possible causes for species belonging to the dark diversity are manifold and include, but are not limited to, lower-level ecological filters involving metapopulation and metacommunity dynamics (cf. dispersal limitations and habitat fragmentation, Tilman, 1997; Fahrig, 2003) or complex biotic interactions (e.g. competition, parasitism, mutualism and symbiotic phenomena like mycorrhiza, Grime, 1979; Torrez *et al.*, 2016). To our knowledge, the likelihood of individual species to belong to the dark diversity has never been studied and little is known about the characteristics of species having a higher chance to belong to the dark diversity pool than others (but see Riibak *et al.*, 2015).

Recently, the dark diversity concept has been proposed as a new facet of biodiversity for gaining useful knowledge for restoration and conservation issues (Pärtel, Szava-Kovats & Zobel, 2011). However, till now only few studies have been inspired by this (Gijbels, Adriaens & Honnay, 2012; Yoshioka *et al.*, 2014; Riibak *et al.*, 2015; Ronk, Szava-Kovats & Pärtel, 2015). Indeed, these have been successful albeit suffered from various weaknesses preventing a more general application: (1) spatial extent or resolution was not directly relevant for practical nature conservation and restoration planning (Riibak *et al.*, 2015; Ronk, Szava-Kovats & Partel, 2015), (2) scope was only a single habitat or ecosystem (Yoshioka *et al.*, 2014; Riibak *et al.*, 2015) or (3) only a few species were involved (Gijbels, Adriaens & Honnay, 2012). Although the restoration potential of a given site may depend on a number of site-specific factors, like fragmentation and biotic interactions within the focal community, searching across multiple sites and habitats for common traits among typical dark diversity species may clarify potential drivers of the observed site-specific species distribution patterns. Hence, studying the features of species that belong to the dark diversity more often than others could prove to be an important key for successful practical application of dark diversity in restoration and conservation.

Clearly, the delimitation of the regional species pool can impact the assessment of dark diversity massively. Consequently, this issue has been one of the major concerns of the original dark diversity approach: if habitats are delimited rather categorically as suggested in the original concept paper (Pärtel, Szava-Kovats & Zobel, 2011) the actual natural environmental gradients are ignored (Mokany & Paini, 2011). For plants, regional species pools have been successfully identified using the species indicator values presented by Ellenberg *et al.* (2001) for Central European plant species (Pärtel *et al.*, 1996). Ellenberg's indicator values (EIVs) represent European plant species' preferred position along various environmental gradients and are often used in vegetation studies (Ellenberg *et al.*, 2001; Diekmann, 2003; Lenoir *et al.*, 2010; Moeslund *et al.*, 2013). Recently, Ewald (2002) proposed a probabilistic procedure (Beals' index) to estimate regional species pools based on co-occurrence patterns among species. This approach is currently known to yield the most realistic estimates of the regional species pool (Lewis, Szava-Kovats & Pärtel, 2016).

Restoration, conservation and nature management typically take place at relatively fine spatial resolutions. Obviously, considering dark diversity at a resolution relevant for conservation and restoration management is likely to inflate the dark diversity simply because smaller areas support fewer species all else being equal (McArthur & Wilson, 1967). For this reason, comparing dark diversity from areas of different sizes where co-occurrence data were collected in a different manner is not meaningful without accounting for these effects. To ensure a reliable assessment of dark diversity, co-occurrence data collected at fine spatial resolution across large spatial extent and in a systematic manner—like in national biodiversity inventories (e.g. Fredshavn, Nygaard & Ejrnas, 2009) − is needed.

Here we present the first national assessment of the characteristics of typical dark diversity plants at a spatial resolution relevant to conservation and restoration management, covering multiple open terrestrial habitats. Using a large national plant dataset with high spatial accuracy and a combination of EIVs, Grime's plant strategies, mycorrhizal information and dispersal distance calculations, we address the following specific study questions: (1) does North-European plant species differ in how often they occur in the dark diversity pool (species’ likelihood to belong to dark diversity)? (2) If so, which plant traits or ecological characteristics explain this pattern the best? Finally, we discuss the causal mechanisms most likely involved and how our findings may aid effective planning and management initiatives and promote the practical application of the dark diversity concept within conservation and restoration.

## Methods

### Vegetation data

Data on the distribution of vascular plants in Denmark was obtained from municipalities’ vegetation inventory of natural habitat types (Fredshavn, Nygaard & Ejrnas, 2009). We used observations from 5-m radius circular plots laid out to capture the typical flora of a particular site in question (typically one plot per site). The sites are 5 ha on average (ranging between 0.003–900 ha), distributed throughout most of Denmark (Fig. 1) and cover freshwater meadows, salt meadows, heathlands, bogs, moors, fens, grasslands and vegetated dunes (i.e. open habitats). The dataset was extracted 6 October 2014. We used data from 2004–2014 encompassing 236,923 records from 15,160 plots involving 564 plant species after application of the filters described in the following. We only considered observations at the species level and excluded all neophytes (Appendix tables 6-8 in Buchwald *et al.*, 2013), i.e. species that are not considered a natural part of the vegetation given their history and dispersal ability), shrubs, trees and submersed aquatic species. To ensure meaningful calculations of the regional species pool (see below) only plots with more than five plant species records were used.

**Figure 1.**
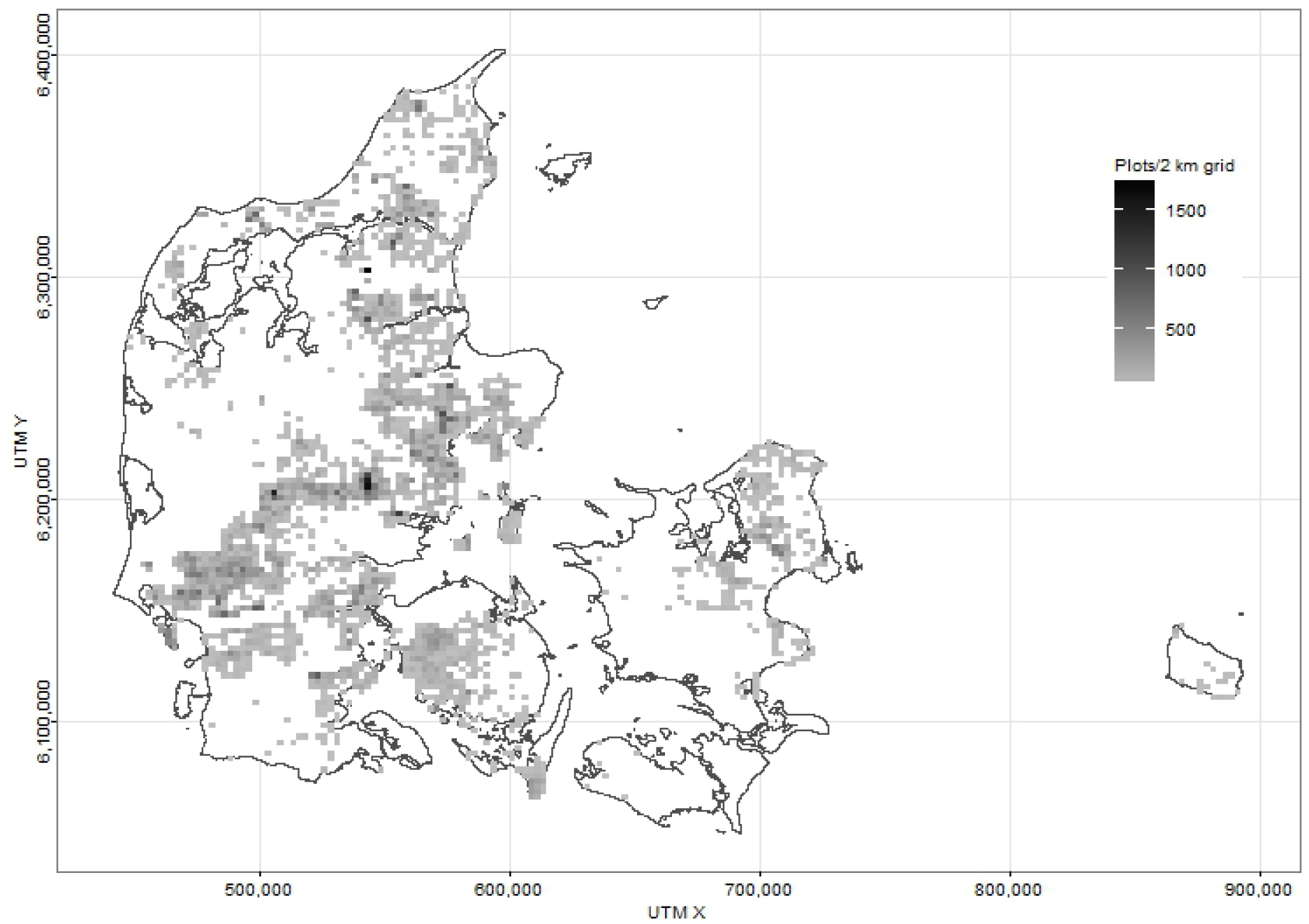
The density of the 15.160 vegetation plots used for this study

### Regional species pool and dark diversity

To yield the best estimates of dark diversity, we used Beals’ index (Beals, 1984) to assess the regional species pool for each plot as recommended by Lewis, Szava-Kovats & Pärtel (2016). For each plot, Beals' index represents the probability that a focal species will occur within that plot given the assemblage of species co-occurring there (McCune, 1994). See details in Münzbergova & Herben (2004). Initially, a presence/absence matrix with all combinations of plot and species was calculated. Based on this matrix, we calculated Beals’ index for each species in each plot excluding the focal species from the calculations (as recommended by Oksanen *et al.*, 2015) (Fig. 2). We used the “beals()” function in the “vegan” package (Oksanen *et al.*, 2015). The threshold for including a species in the regional species pool was defined as the 5^th^ percentile of the Beals’ index value for the species following Gijbels, Adriaens & Honnay (2012) as well as Ronk, Szava-Kovats & Pärtel (2015). Additionally, we only considered data for plots having Beals’ index values above that of the lowest value where the species was indeed present. For every plot the dark diversity was composed by all species in the regional pool excluding those that were actually present (Partel, Szava-Kovats & Zobel, 2011) (Fig. 2).

**Figure 2.**
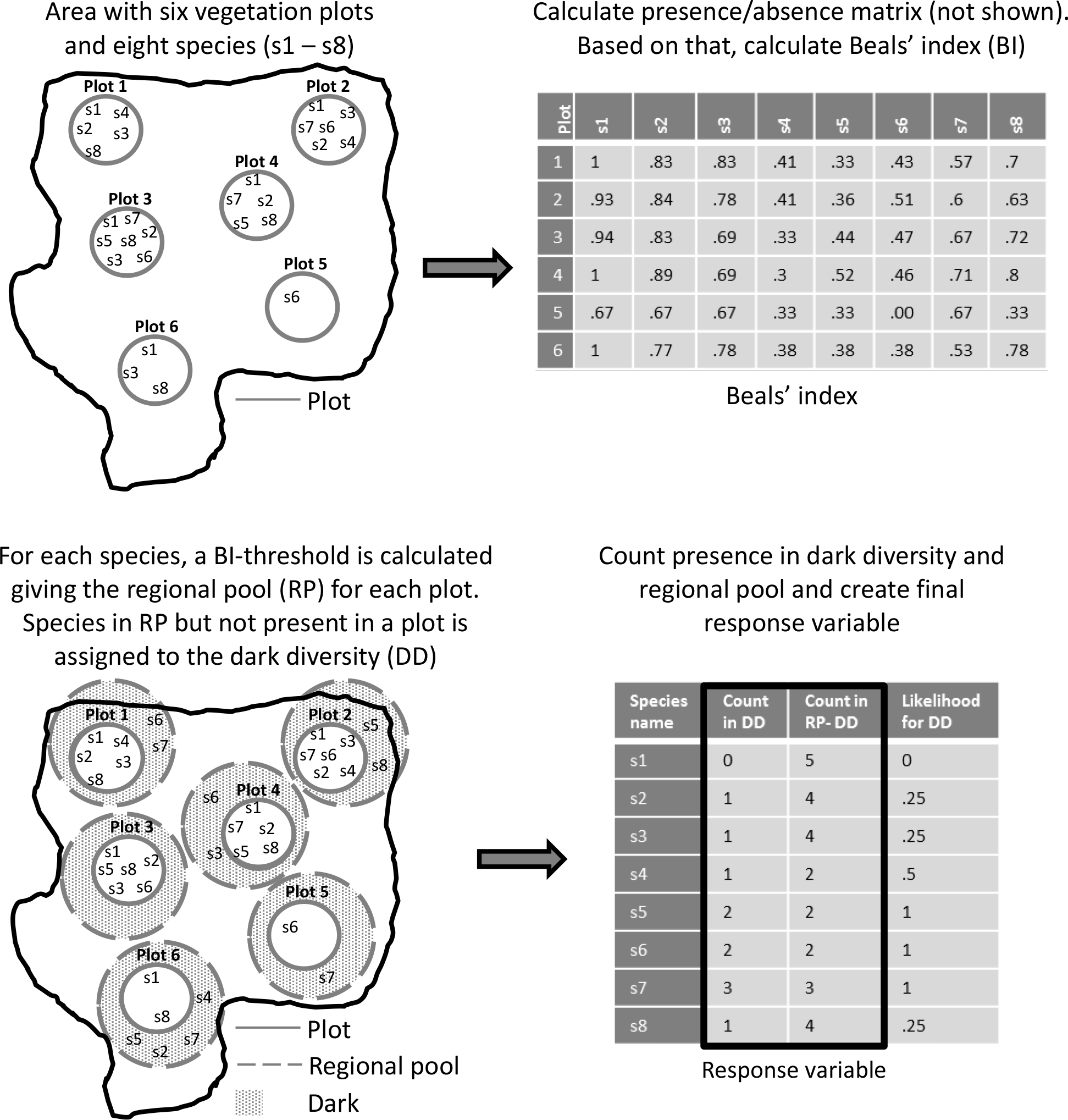
The steps taken from vegetation plot data to response variable. A fictive area with eight fictive species is used for illustration purpose

### Response variable

As a response variable, we computed the species’ likelihood to belong to dark diversity (cf. the ratio of the number of occurrences in the dark diversity pool divided by the number of occurrences in the regional species pool) (Figs 2 & 3) for each of the 564 plant species used in our analyses.

**Figure 3.**
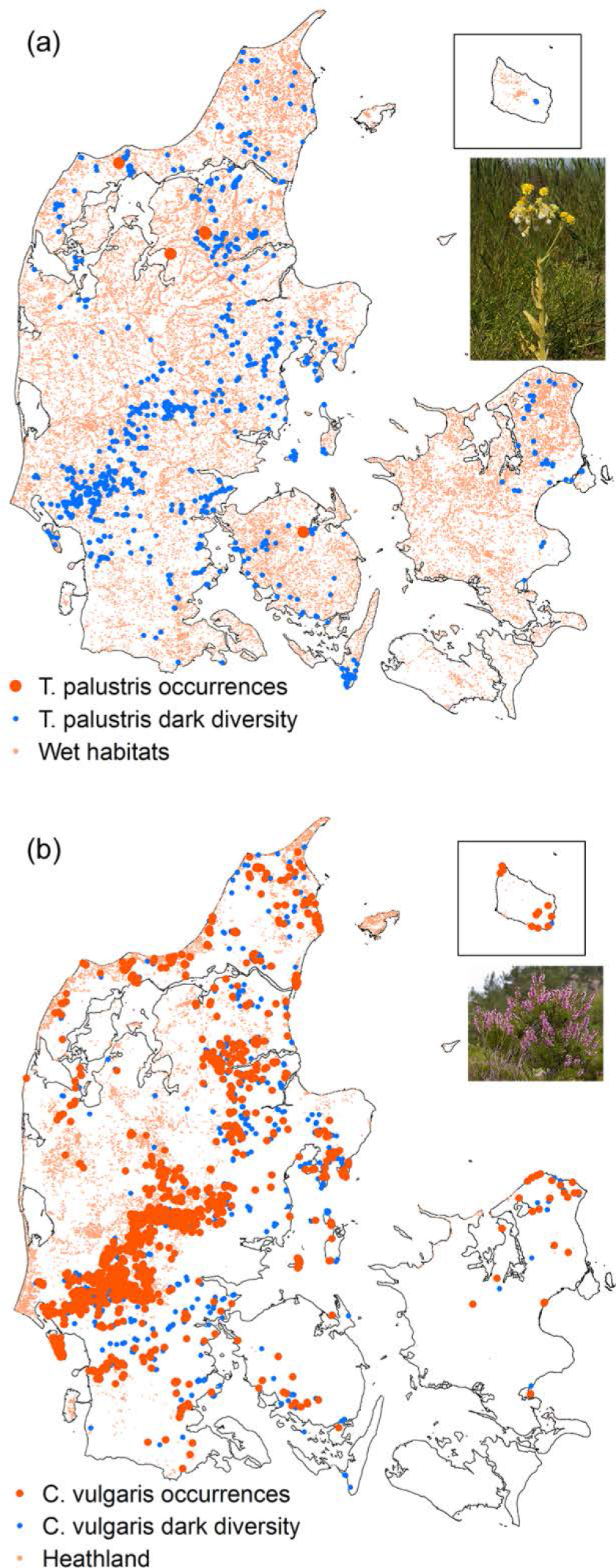
The distribution of occurrences and dark diversity locations for (a) a species often found in dark diversity (Tephroseris palustris) and (b) a species relatively rarely found in the dark diversity in this study (Calluna vulgaris). For reference, known wet habitats and heathlands are shown. Map inserts: island of Bornholm (Fig. 1)

### Species traits and characteristics (explanatory variables)

#### Ellenberg's indicator values

We used Ellenberg's indicator values (EIVs) (Ellenberg *et al.*, 2001) adjusted to British conditions (Hill *et al.*, 1999) as these adjusted values are thought to best match the Danish flora (Moeslund *et al.*, 2013).Variation in temperature (EIV_T_) and continentality (EIV_K_) in Denmark is negligible and salinity (EIV_S_) is only relevant in coastal environments. Consequently, we only considered EIVs for: (1) soil moisture (EIV_F_), (2) soil pH (EIV_r_), (3) soil nutrient status (EIV_N_) and (4) ambient light (EIV_L_) (Table S1). EIVs for soil nutrient status and pH are typically highly correlated (Diekmann & Falkengren-Grerup, 1998; Seidling, 2005); hence we calculated a nutrient/pH-ratio (EIV_N_/_R_) as an alternative variable based on the two corresponding EIVs. This variable represents the species’ preference for nutrient availability (see for example Andersen *et al.*, 2013).

#### Grime plant strategies

The main plant strategies presented by Grime (1979) enables scientists to distinguish between plants adapted to competitive (C-species), stressful (S-species) or ruderal (R-species) environments. Although plants can harbour any combination of these three strategies they are in their extreme forms mutually exclusive (Grime, 1979). For this study, we obtained the strategies for each plant species from the BiolFlor database (Kuhn, Durka & Klotz, 2004). Grime plant strategy data was available for all species included here (Table S1). Following Ejrnas & Bruun (2000), we represented the degree to which a plant is adapted to a given strategy as values ranging from 1-12 for each of the three strategies, however restraining their sum to 12.

#### Mycorrhiza data

We used data on both mycorrhizal infection percentage (0 to 100%) and dependence (i.e., a factor variable with two levels: obligately vs. not obligately mycorrhizal). Data on mycorrhizal infection was retrieved from Akhmetzhanova (2012) and data on mycorrhizal dependence was taken from MycoFlor (Hempel *et al.*, 2013). These data were available for 33 % and 82 %, respectively, of the plant species involved in this study (Table S1).

#### Plant functional traits

The trait data has two purposes: (1) it is used for modelling dispersal distance (see below) and (2) seed mass is used as an explanatory variable representing an alternative measure of dispersal distance as well as the plants ability to establish at new sites. We obtained seed mass (SM), dispersal syndrome (DS), releasing height (RH), terminal velocity (TV) and growth form (GF) data from the LEDA and BiolFlor databases (Kuhn, Durka & Klotz, 2004; Kleyer *et al.*, 2008). Where multiple records of SM, RH and TH were available for the same species, the mean value was used. Missing data on DS and GF was taken from Hansen (1996) making these two traits available for all species (94 % and 77 % of the species respectively were covered by the aforementioned databases). Data on SM, RH and TV was available for 89 %, 91 % and 68 % of our study species, respectively.

Subsequently, we calculated maximum dispersal distance (MDD) using the “dispeRsal()” function (Tamme *et al.*, 2014). This function calculates MDD using plant traits and taxonomy. For 70% of the species, MDD was calculated with a model including DS, GF and TV. For the remaining 30% of the species MDD was calculated using simpler models following the hierarchy of best predictive performance given in Tamme *et al.* (2014). For species with multiple different entries of a trait (e.g. DS) we calculated the mean of the predicted MDDs. Table S1 lists the MDDs calculated for each species.

### Data analysis

We used binomial generalised linear models (GLMs) for proportion data to explore the relationship between the plants' likelihood of being part of the dark diversity (binomial response variable; no. of times in dark diversity/no. of times in the regional species pool) and the 13 explanatory variables listed in Table 1. All variables were tested for multicollinearity (see Figure S1). The EIVs for nutrients and pH both showed a high degree of multicollinearity (tolerances below 0.15, Quinn & Keough, 2002) and therefore we decided to use the nutrient/pH-ratio (described above) instead to represent the plants’ nutrient preferences (Table 1).

**Table 1.** An overview of the included explanatory variables with a brief description of each variable. For abbreviations see Table 2

| Explanatory variable | Short description | Variable nature | Relevant literature |
| --- | --- | --- | --- |
| EIV<sub>L</sub> | Plants' preference for light level | Ordinal (1–9) | (Ellenberg et al. 2001) |
| EIV<sub>F</sub> | Plants' preference for soil moisture | Ordinal (1–12) | (Ellenberg et al. 2001) |
| EIV<sub>N/R</sub> | Soil nutrients and pH is usually positively correlated. To take this into account this ratio is used to represent nutrient enrichment | Continuous (0.13 – 2.00) | (Ellenberg et al. 2001, Andersen et al. 2013) |
| Grime C | Plants' competitive ability | Ordinal (1–12) | (Grime 1979) |
| Grime S | Plants' stress tolerance (e.g., tolerance towards grazing, cutting, drought, soil salinity etc.) | Ordinal (1–12) | (Grime 1979) |
| Grime R | Plants' ruderality (tendency to occur in ruderal areas) | Ordinal (1–12) | (Grime 1979) |
| Mean mycorrhiza infection % | The mean infection percentage of the plants' roots by mycorrhiza | Continuous (1–100 %) | (Akhmetzhanova et al. 2012) |
| Mycorrhizal dependence | Whether (1) or not (0) the plants are obligately mycorrhizal. Binary variable | Nominal (0 vs. 1) | (Hempel et al. 2013) |
| Log(Seed mass) | The logarithm to the mean seed mass | Continuous (-2.52 – 2.13) | (Kühn et al. 2004, Riibak et al. 2014) |
| Log(Max. dispersal distance) | The logarithm to the maximum dispersal distance based on plant traits and taxonomy | Continuous (-1.13 – 10.72) | (Tamme et al. 2014, Riibak et al. 2014) |

**Table 2.**
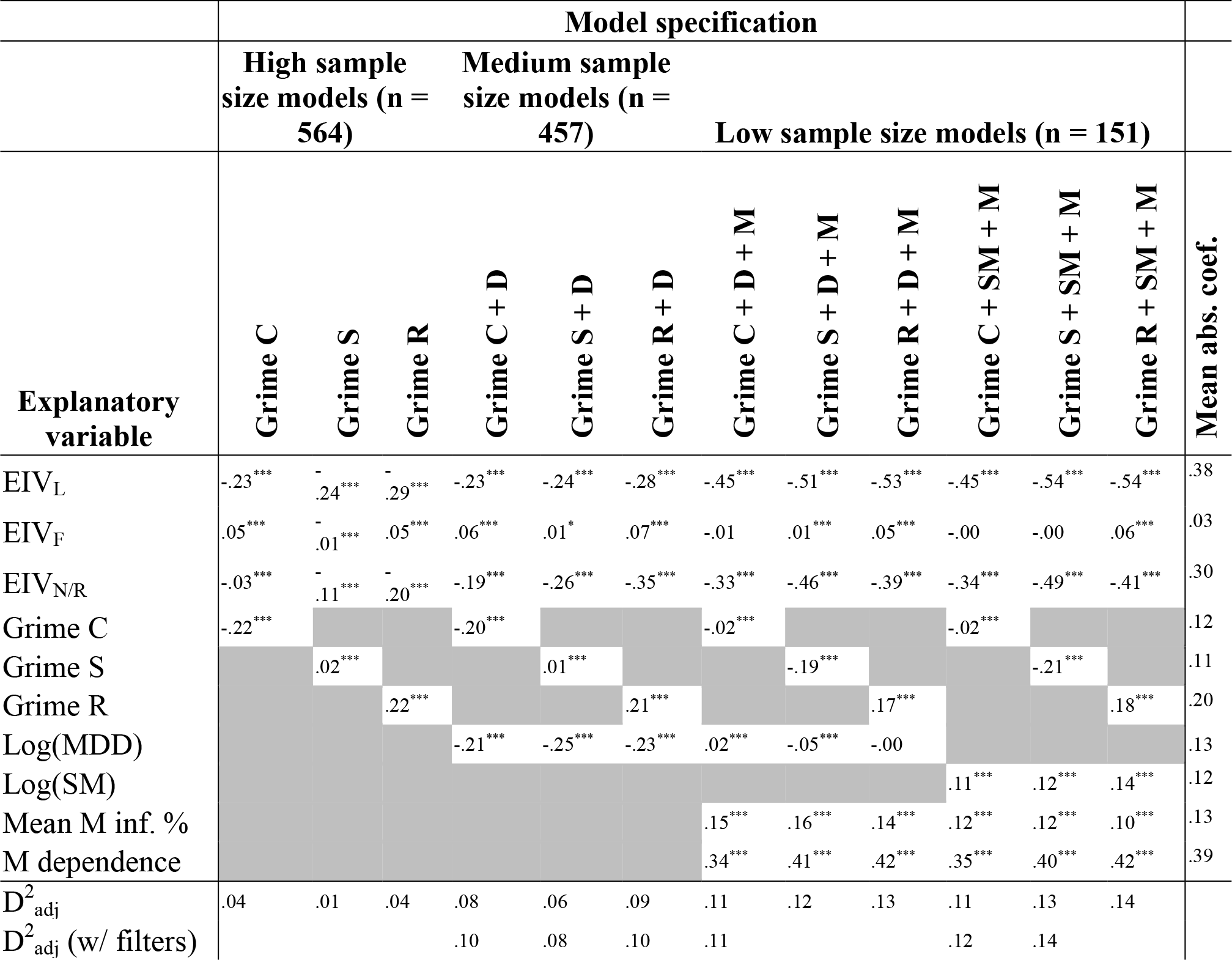
Standardised coefficients (their rank based on the numeric size of the standard coefficient is given in parentheses) of each of the 12 candidate models relating the probability for plant species of being part of the dark diversity to the explanatory factors used in this study. Results are presented both for the high-sample size models, medium sample size models and the low sample-size models (See details in Table S2). Grey cells mark explanatory variables that were not included in the model. For each model the deviance explained–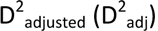–is shown together with the 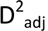 for models after adding the necessary number of filters (also shown) to remove phylogenetic autocorrelation (see methods). For each explanatory variable the absolute (abs.) mean standardised coefficient (coef.) is given to represent its overall relative importance. Abbreviations: C: competition, S: stress, R: ruderality, Disp.: dispersal, Dist.: distance, SM: seed mass, EIV: Ellenberg Indicator for light (L), moisture (F), nutrients (N) and reaction (R), M.: mycorrhiza

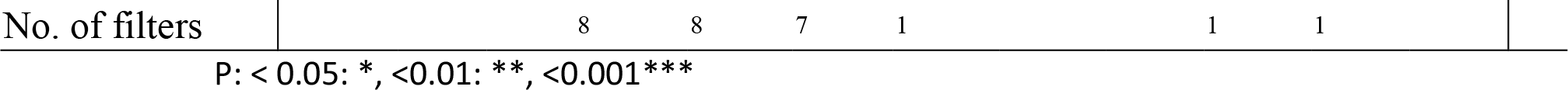

Not all traits and characteristics were available for every species in the dataset (see method sections above). In order to maximize the sample size we initially ran model selection (see section below) on models including only the explanatory variables which were available for the majority of the species (n = 564, Table S2). These models are referred to as *high sample size models* in the following. Being available for a lower number of species (n = 457) dispersal distance was subsequently added to the models subject to model selection *(medium sample size* models). Finally, we ran model selection on models with all explanatory variables listed in Table 1 (n = 151, *low sample size* models). Since the three Grime-based variables were highly dependent on each other, only one Grime variable was included in a model at a time. Also, since the calculations of maximum dispersal distance involved seed mass these two variables never occurred simultaneously in any of the models to avoid redundancy. Following the above-mentioned constraints, we built three sets of candidate models and used Akaike’s Information Criteria (AIC, Aikaike, 1974) to select the best model within each sample size group following best practice as recommended in Burnham & Anderson (2002) (for details on model selection and setup, see Table S2).

A typical issue in the investigation of plant species traits and characteristics is phylogenetic autocorrelation, i.e. the fact that closely related species also tend to be more similar in traits and characteristics (Gittleman & Kot, 1990). Accordingly, we checked all models’ residuals for phylogenetic autocorrelation as described below. We used the Daphne phylogenetic tree for the European flora (Durka & Michalski, 2012) and followed the vignette by Paradis (2015) to calculate Moran's I of each model’s residuals using the reciprocal phylogenetic distances between species. This computation was performed using the “Moran.I()” function in the “ape” package (Paradis, Claude & Strimmer, 2004). Six models showed significant phylogenetic autocorrelation. To account for this we constructed phylogenetic eigenvector filters following best practise (Borcard & Legendre, 2002; Diniz-Filho & Bini, 2005). Filters that explained significant variation in the best models’ residuals were added successively to these best models in addition to the explanatory variables. After each addition of a filter, the model residuals were checked for autocorrelation. When Moran’s I showed no significant autocorrelation the process stopped. Significance and effect sizes of explanatory variables in the models were reassessed but no further model selection was performed.

All analyses were conducted using R (R Core Team, 2015).

## Results

On average, species were part of the dark diversity in 88.6 % of the plots for which the species was indeed in the regional pool (Table S1).

We found phylogenetic autocorrelation in the residuals of six models (Moran's I test, P < 0.05). The addition of 1 to 8 phylogenetic filters successfully removed autocorrelation and caused no notable shifts in effects sizes or significance (Table 2, Table S3).

The goodness of fit for our models was up to 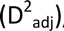, with the more complex low sample size models–including potential maximum dispersal rate, average seed mass and mycorrhizal information–having the best fits (Table 2). Our models suggest that the factors best explaining the plants’ likelihood of being in the dark diversity are (mentioned in the order of importance): mycorrhizal dependence, preference for light and nutrients, ruderality, maximum dispersal distance, seed mass, mycorrhizal infection percentage, stress-tolerance, competitive ability and preference for soil moisture (Table 2). We found strong indications that obligate mycorrhizal plants are more often part of the dark diversity than plants not depending on mycorrhiza (Fig. 4). This finding was supported by the fact that species with higher dark diversity likelihood had a higher degree of infection by mycorrhiza (Table 2, Fig. 4). In addition, plants more frequently in the dark diversity were adapted to thrive under low nutrient availability and low-light (Fig. 4), had poor dispersal abilities and heavier seeds. In terms of Grime strategies, the high dark-diversity-likelihood species were more ruderal, less competitive and less stress-tolerant (Table 2).

**Figure 4.**
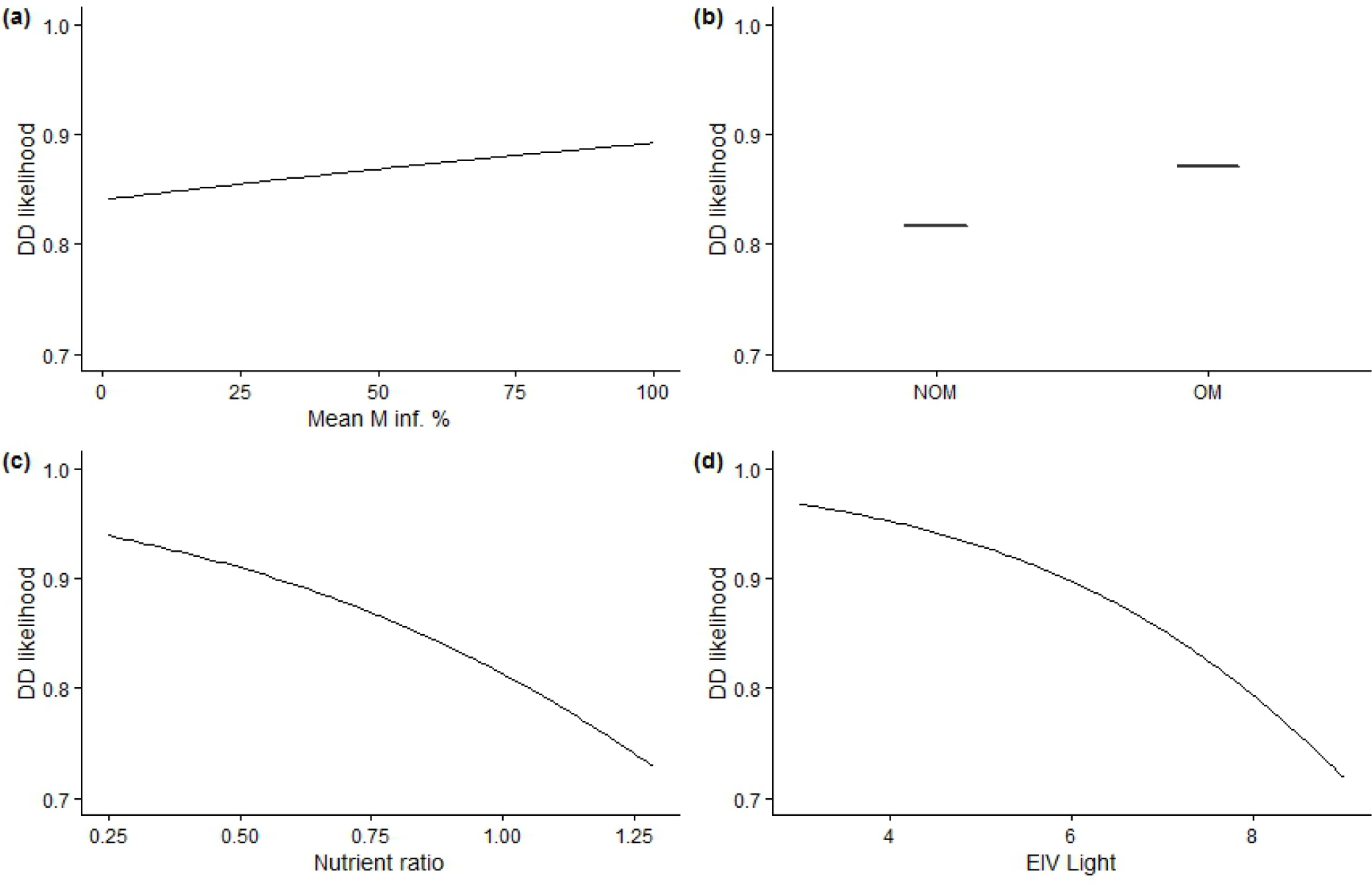
Predictions of the relationship between the dark diversity likelihood of a plant and (a) mean mycorrhizal infection percentage, (b) mycorrhizal dependence, (c) nutrient/reaction-ratio (nutrient ratio) based on Ellenberg Indicator Values (EIV) and (d) EIV for light. The predictions shown are based on the *Grime R + SM + M* model having the highest goodness of fit (Table 2). Predictive variables not in question were held constant during the predictions shown. DD: dark diversity, NOM: not obligately mycorrhizal, OM: obligately mycorrhizal, M: mycorrhizal, inf.: infection

## Discussion

### Differences in species’ dark diversity likelihood

Intuitively, typical dark diversity species are rarer than their habitat would suggest, for example rare species with a fairly common habitat (Fig. 3). A species such as *Tephroseris palustris* (L.) Rchb. was often found in the dark diversity (Fig. 3a). This species is rare in Denmark although it has a wide distribution in Northern Europe including Scandinavia, the Baltic region and all the way to arctic Russia (Kochjarová, 2006). It is extinct in Great Britain, the Czech Republic, Hungary, Slovakia and Romania (Kochjarová, 2006) and critically endangered in Sweden (Olsson & Tyler, 2001). However, it tends to colonize bare mud along pond margins and is even known from recycling depots (Frederiksen, Rasmussen & Seberg, 2006)–habitats that are far from rare in Denmark. Hence, habitat availability alone cannot explain its rarity. This was also the case for *Leontodon hispidus* L., *Campanula persicifolia* L., *Vicia tetrasperma* (L.) Schreb. and several others (Table S1). On the other hand, species that occurred less often in the dark diversity were in many cases species that are common within their habitats, such as *Calluna vulgaris* (L.) Hull (common in heathlands, Fig. 3) and *Tripolium vulgare* Nees (common in salt meadows) or alternatively they were common generalists like *Achillea millefolium* L., *Festuca rubra* L., *Urtica dioica* L. etc. (Frederiksen, Rasmussen & Seberg, 2006).

### The relative importance of explanatory factors and their likely causal mechanisms

Apart from stochasticity (Hubbell, 2001), there are generally three factors important for species’ dark diversity likelihood: (1) dispersal ability, (2) establishment success and (3) persistence in a given habitat. The explanatory factors tested in this study all fall within these three categories. Factors primarily involved in establishment and persistence were the most important ones in our study overall: mycorrhizal dependence and the plants’ preference for available nutrients and light. Dispersal related factors such as the plants’ maximum dispersal distance and the seed mass were also important, with the latter playing a role for both dispersal and establishment (see below).

#### Establishment and persistence

A key ability in plants’ life cycle is their establishment once the seed has settled at a given site (Muller-Landau *et al.*, 2002). Many factors such as seed herbivory, seed resistance to pathogens, stress-events like drought or flooding and the seeds’ endosperm resources can affect this ability (Maun, 1994; Muller-Landau *et al.*, 2002; Moles & Westoby, 2004). Another imperative aptitude for establishment of some plants is the development of mycorrhizae (Akhmetzhanova *et al.*, 2012; Gijbels, Adriaens & Honnay, 2012; Hempel *et al.*, 2013). Additionally, mycorrhizae have been shown to be important not only in the establishment phase but also for the persistence of plant species and consequently for the local plant community composition (Hartnett & Wilson, 1999). Here we demonstrated that plants depending on mycorrhiza and plants that require a high degree of mycorrhizal infection had higher dark diversity likelihood. This result is not surprising given the importance of mycorrhiza for plant establishment and persistence.

Competition for resources is an important phenomenon shaping the local structure, composition and richness of plant communities throughout the world (Tilman, 1994; Tilman, 1997; McKane *et al.*, 2002; Moeslund *et al.*, 2013). In the current study, we showed that plants with high dark diversity likelihood preferred low levels of nutrients. In most of Northern Europe and particularly in Denmark the landscape is relatively nutrient rich. In such a setting, plant species like *Urtica dioica* L., *Epilobium hirsutum* L. and *Cirsium arvense* (L.) Scop. adapted to benefit from high nutrient availability will be strong competitors (Grime, 1979; Hill *et al.*, 1999; Ellenberg *et al.*, 2001) and therefore have lower dark diversity likelihood. Plants with competitive advantages (e.g., regarding resources) will tend to appear more often in suitable habitats than species being less strong competitors in these habitats.

In our study demonstrated that shade tolerant plants were more likely to be part of the dark diversity. The possible explanation for this finding is probably multifaceted. Plants that are more or less shade tolerant may have competitive disadvantages in open landscapes (recall that this study concerns only open habitats). Also, the fact that the landscape was prehistorically (around 5000 BC) almost completely forested (Fritzboger & Odgaard, 2010) can explain this observation. Indeed, shade tolerant species likely to occur dormantly in the soil seed bank may have been present and more frequent across Denmark during the past, when forests were more dominant, and may return (cf. memory effect due to land-use legacy) if land-use returns to forest habitats. A third explanation could be that since forests are not included in this study some shade tolerant species may have higher dark diversity likelihood simply because they prefer forest environments. While there are indeed examples of such “forest-species” that occasionally occur in open habitats in our dataset (e.g. *Galium odoratum* (L.) Scop. and *Allium ursinum* L.), there are also several examples of shade tolerant species often belonging to the dark diversity albeit in reality observed almost as often in open landscapes as in forests: e.g. *Maianthemum bifolium* (L.) F.W.Schmidt and *Trientalis europaea* L. (EIV_l_ = 3 and 5 respectively, both widespread in grasslands, meadows, heathlands, forests etc.) (Mossberg & Stenberg, 2005).

In many environments, stress tolerance is a key factor shaping local plant diversity (Osmond *et al.*, 1987; Maun, 1994; Ejrnas & Bruun, 2000; Moeslund *et al.*, 2011). Recently, researchers showed that stress-tolerance was among the most important determinants of plant dark diversity in North-Eastern European dry calcareous grasslands (Riibak *et al.*, 2015). They suggested that in the driest grasslands stress-tolerant species are more likely to thrive and therefore less likely to be part of the dark diversity. For our study this explanation could also be true. Many of the habitats we included are stressful environments either because they are relatively dry (grasslands, heathlands), waterlogged (fens, bogs, meadows) or saline (salt meadows). However, another equally plausible explanation could be that ruderal species occurred more often in the dark diversity than non-ruderals and the plants’ ruderality was significantly negatively related to the plants’ stress-tolerance. From the fact that highly ruderal species cannot be highly stress-tolerant at the same time (Grime, 1979) it follows that the likelihood of ending up in the dark diversity could actually be related to ruderality, not stress-tolerance.

#### Dispersal

We showed that species with higher dark diversity likelihood were also generally poorer dispersers. This result was supported by Riibak (2015) for dry calcareous grasslands. It is intuitively meaningful and a well-known fact that species with poor dispersal abilities also have a lower probability of dispersal to new suitable sites and to recolonize sites where these species have earlier gone extinct (Tilman, 1997; Cain, Milligan & Strand, 2000; Myers & Harms, 2009; Torrez *et al.*, 2016). This explanation is corroborated by the result that species with a higher seed mass was also more often part of the dark diversity. Heavy seeds are more unlikely to spread long distances and thereby reach suitable habitats (Marteinsdóttir & Eriksson, 2013). Also, heavy-seed species typically produce less seeds lowering the probability that one of them will eventually arrive at a suitable site (Cornelissen *et al.*, 2003; Riibak *et al.*, 2015).

### Explaining dark diversity likelihood

As expected–given their importance for plant species establishment, dispersal or persistence–the factors included here were able to significantly explain plant species’ dark diversity likelihood. The goodness of fit 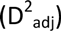 we obtained implies that even though most of the explanatory factors were important for plants’ dark diversity likelihood other factors not tested here may also be involved. Site conditions such as habitat fragmentation and reduced habitat patch sizes may be important issues that can explain why many species are missing in suitable places (Fahrig, 2003). However, the plants’ ability to survive in a fragmented small-habitat-patch landscape is not trivial to measure and attempts to directly relate plant traits to this ability are therefore rare (Dupre & Ehrlen, 2002; May *et al.*, 2013). We attempted to account for this by including dispersal distance and seed mass, as suggested by May *et al.* (2013) and Grime’s plant strategies, which relate to seed number, life span, seed bank strategy and pollination strategy (Grime, 1979; Rees, 1994; Sera & Sery, 2004; Pierce *et al.*, 2014), which again relate to species’ probability to go extinct in a suitable habitat patch (Eriksson, 1996; Dupre & Ehrlen, 2002; Gijbels, Adriaens & Honnay, 2012). On the other hand, researchers have recently found that while seed production traits are integral to Grime’s plant strategies this relationship is not straight-forward (Pierce *et al.*, 2014). This might also be true for relationships between Grime’s plant strategies and factors such as life span and pollination strategy and therefore future studies may obtain a higher goodness of fit by analysing the individual traits instead of proxies for these. Furthermore, phenotypic plasticity and general susceptibility to pathogens are likely to play a role for the persistence of plants (Augspurger & Kelly, 1984; Burdon, Thrall & Ericson, 2006; Reed *et al.*, 2010).

### Management implications

In the EU, part of the biodiversity strategy is to halt biodiversity loss by 2020. This goal requires both researchers and managers to make the most of the limited funds available. Restoring or conserving ecosystems or habitats is often focused on re-establishing or improving environmental conditions and then hoping that biodiversity will respond positively. Tools that improve our understanding of the causal mechanisms behind which species remain absent and why despite seemingly suitable conditions is extremely useful and could be highly beneficial for restoration and conservation efforts.

For conservation and restoration our findings underpin the importance of assessing the funga’s ability to sustain the flora of concern at a given site. Inoculation with certain fungi could aid to restore plant communities (Torrez *et al.*, 2016). However, great care needs to be taken during this proces with the best results probably obtained by adding a diverse and locally adapted mycorrhizal community (Klironomos, 2003).

The fact that dispersal limitation is an important factor for species’ dark diversity likelihood strongly suggests that space and time are imperative factors in the planning and management of nature. Given enough time and suitable corridors for dispersal, even poor dispersers will eventually reach suitable but distant habitats. This finding also highlights the need to accommodate this by considering assisted migration (Seddon, 2010).

Also, our study suggest focusing on creating opportunities for ruderal species in restoration and conservation projects; e.g. by making sure that bare soil for seed germination is made available through disturbances such as erosion, fire, trampling by large herbivores etc.

Finally, our results strongly emphasize the importance of focusing on nutrients and light availability in conservation and restoration. Failing to ensure nutrient poor sites in regions with heavy nutrient loads will render a number of species unable to thrive in otherwise suitable areas. Creating more shade in open habitats could potentially aid low-light-adapted species in reappearing from the dark diversity pool.

## Acknowledgements

We are grateful to 15. Juni Fonden for economic support.

### Data accessibility

– Species geography: www.naturdata.dk
– Species traits: LEDA, BiolFlor and MycoFlor databases, Akhmetzhanova *et al.* (2012), TableS1

